## Supplemental Figures for "Distinct cortico-striatal compartments drive competition between adaptive and automatized behavior"

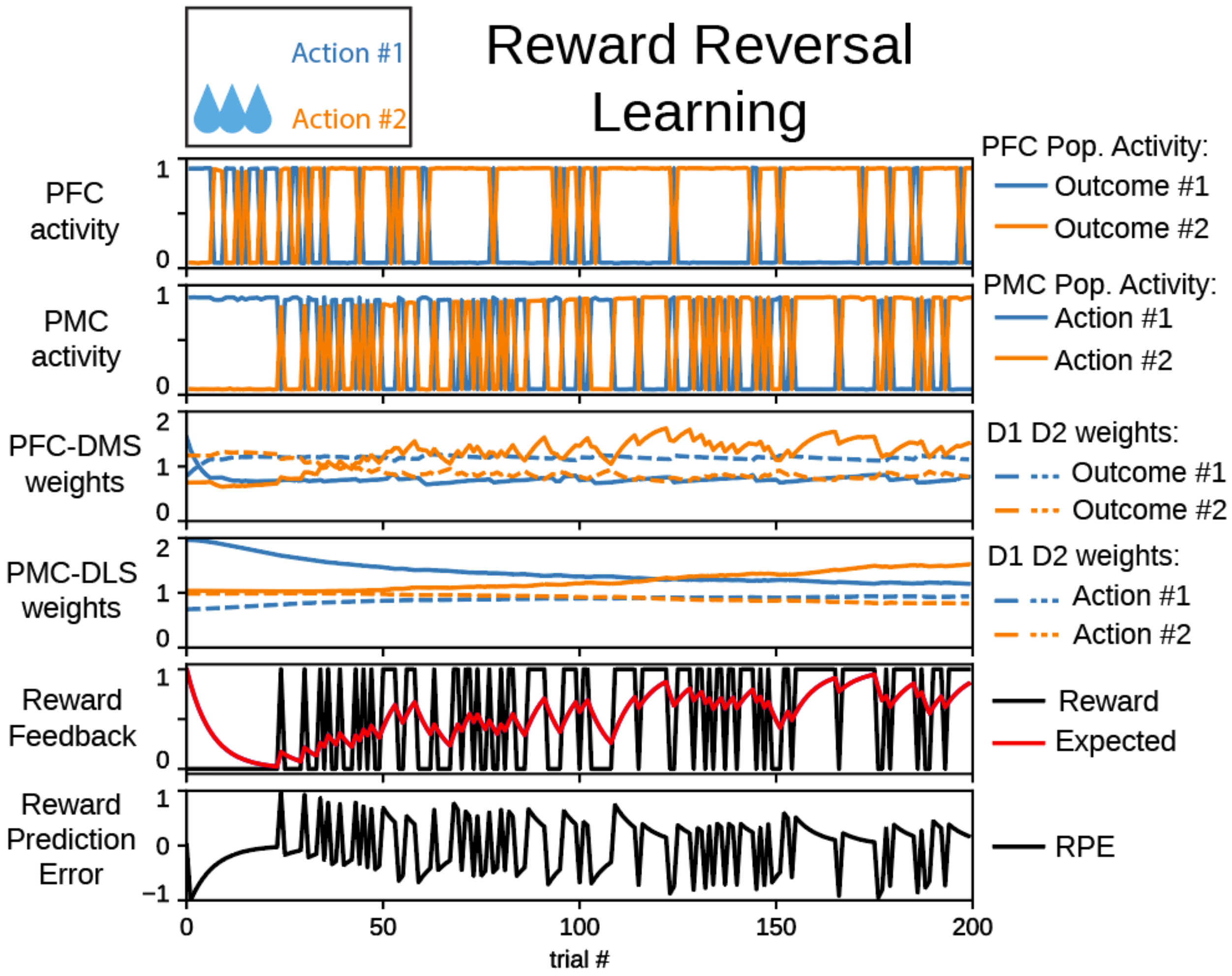


Figure S1. An agent learns to select action #2 in a reward reversal behavioral task following impairment of executive function. In panels depicting cortico-striatal weights, D1 synaptic weights are solid traces and D2 synaptic weights are dashed traces. Blue traces correspond to action #1 and orange traces correspond to action #2.

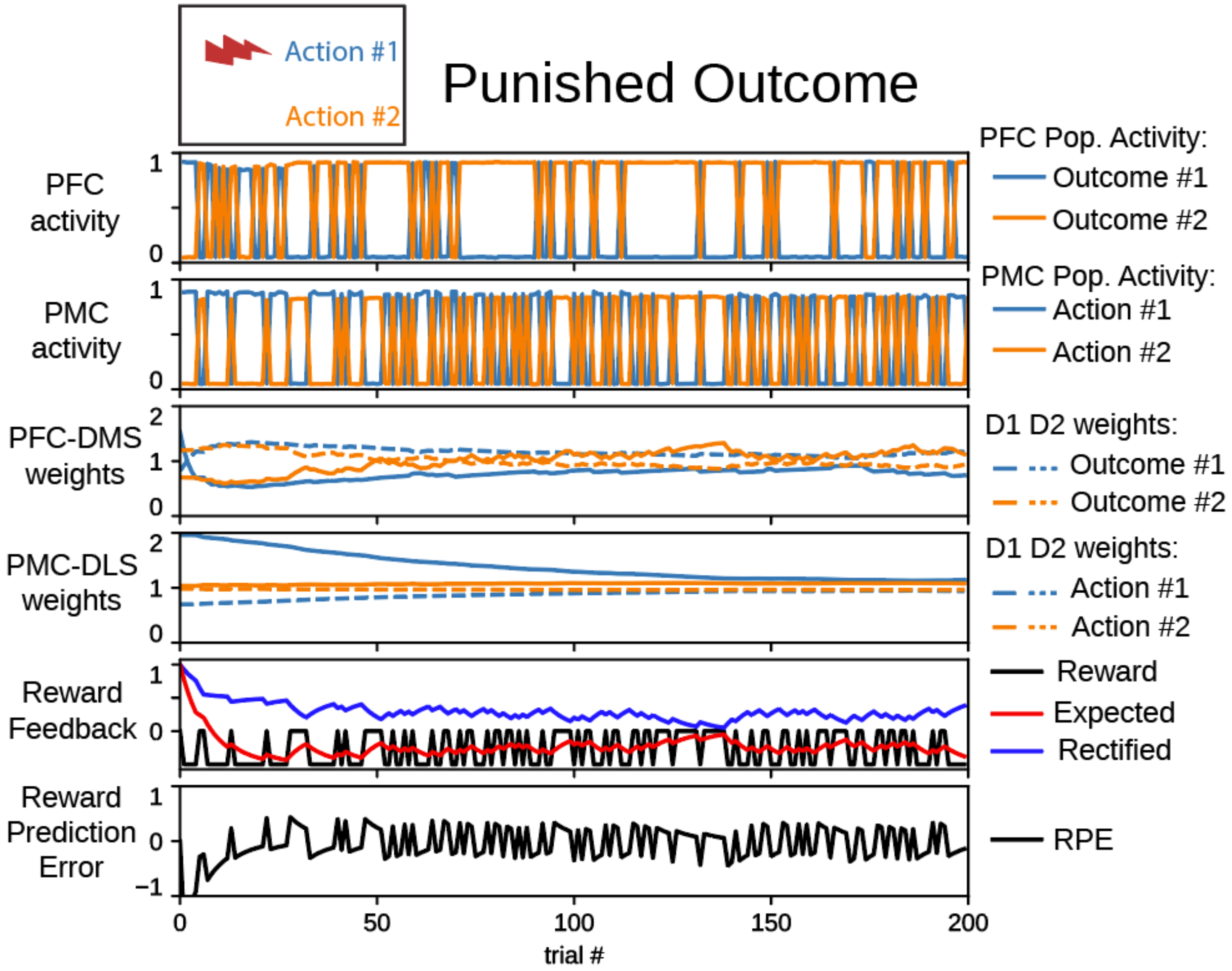


Figure S2. An agent learns to select action #2 in a punished outcome behavioral task following impairment of executive function. In panels depicting cortico-striatal weights, D1 synaptic weights are solid traces and D2 synaptic weights are dashed traces. Blue traces correspond to action #1 and orange traces correspond to action #2.
